# Spontaneous rate of clonal mutations in *Daphnia galeata*

**DOI:** 10.1101/2020.08.31.275495

**Authors:** Markus Pfenninger, Halina Binde Doria, Jana Nickel, Anne Thielsch, Klaus Schwenk, Mathilde Cordellier

## Abstract

Mutations are the ultimate source of heritable variation and therefore the fuel for evolution, but direct estimates exist only for few species. We estimated the spontaneous nucleotide mutation rate among clonal generations in the waterflea *Daphnia galeata* with a short term mutation accumulation approach. Individuals from eighteen mutation accumulation lines over five generations were deep genome sequenced to count de novo mutations that were not present in a pool of F1 individuals, representing the parental genotype. We identified 12 new nucleotide mutations in 90 clonal generational passages. This resulted in an estimated haploid mutation rate of 0.745 x 10^-9^ (95% c.f. 0.39 x 10^-9^ − 1.26 x 10^-9^), which is slightly lower than recent estimates for other *Daphnia* species. We discuss the implications for the population genetics of Cladocerans.

## Introduction

The rate at which spontaneous mutations occur as well as their mutational spectrum influence many important evolutionary parameters and processes. It is relevant for the equilibrium rate of genomic base composition (Hiroshi Akashi and Eyre-Walker 2012), genetic diversity of populations (Johnson and Barton 2005) and the occurrence rate of genetic diseases (Acuna-Hidalgo, et al. 2016). The *de novo* mutation rate determines the possibility (Pfenninger, et al. 2015) and speed of adaptation (Sniegowski and Gerrish 2010) to different environmental conditions. Not the least, knowledge of the mutation rate is essential to estimate effective population size (Charlesworth 2009), population history (Schiffels and Durbin 2014) or divergence times (Ho 2014).

However, direct estimates of the mutation rate exist only for few species because the logistical challenges for such estimations are numerous. Recently, a new approach was introduced that allows an estimation to be made with reasonable effort (Oppold and Pfenninger 2017). Essentially, the approach combines the advantages of mutation accumulation lines with those of the trio approach, while avoiding their respective draw-backs (Oppold and Pfenninger 2017). We adjusted this method here to estimate the clonal mutation rate of the water flea *Daphnia galeata.*

Species of the genus *Daphnia* served since decades as model organisms in ecology, evolution and ecotoxicology (Herrmann, et al. 2018; Miner, et al. 2012; Zhang, et al. 2019). *D. galeata* belongs to the *D. longispina* species complex which dominates the zooplankton of many freshwater lakes in the Holarctic (Ishida and Taylor 2007). The species, like most *Daphnia*, reproduce via cyclic parthenogenesis (Zaffagnini 1987). For most of the time, the species reproduces asexually with a generation time of a few days, while sexual reproduction usually takes only place when environmental conditions deteriorate, usually once or twice a year. The large majority of generational passages are therefore asexual and likely govern the overall rate of mutational change in these species. It was now supplemented with a highly contiguous genome draft (Nickel, et al. in prep.) and other genomic resources (Huylmans, et al. 2016), which allowed the estimation of the clonal mutation rate.

## Material and Methods

### Setting up short term mutation accumulation lines

We used three clonal lines (M5, J2 and LC3.6) to start 24 short term mutation accumulation lines (MAL). These clonal lines were hatched from resting eggs sampled in sediment cores from Müggelsee, Lake Constance (both Germany) and Jordan Reservoir (Czech Republic, see Herrmann, et al. 2018 for details) and maintained in the laboratory since. Details on the laboratory conditions for the general maintenance of *Daphnia* clonal lines can be found in Tams, et al. 2018. In short, single *Daphnia* individuals were cultured in 50 ml artificial *Daphnia* medium (Klüttgen et al. 1994) at 18 +/- 1°C and a light:dark cycle of 16:8 hours. *Daphnia* individuals were fed three times a week with *Acutodesmus obliquus* (1 mg C/ml) and medium was changed weekly.

A single individual from each clone was chosen as F_0_ ancestor for eight MALs for each respective clone. As it is not possible to re-sequence the genome from a single individual, the produced broods 1-3 and 6-11 were raised, pooled and stored for sequencing. This followed the rationale that this ancestor reference pool represents the genotype of the ancestral individual, because mutations occurring in this first generational passage will not dominate the pool but rather appear in singleton reads. The MA-lines were then started from fourth and fifth broods, sisters to the F1 frozen for ancestor reference pool. This proceeding was maintained for the next four generational passages until generation F5. From this generation, all broods (up to sixteen, F6 individuals) were again pooled and used for re-sequencing. A schematic representation of the experimental design can be found in Figure 1.

**Figure 1.**
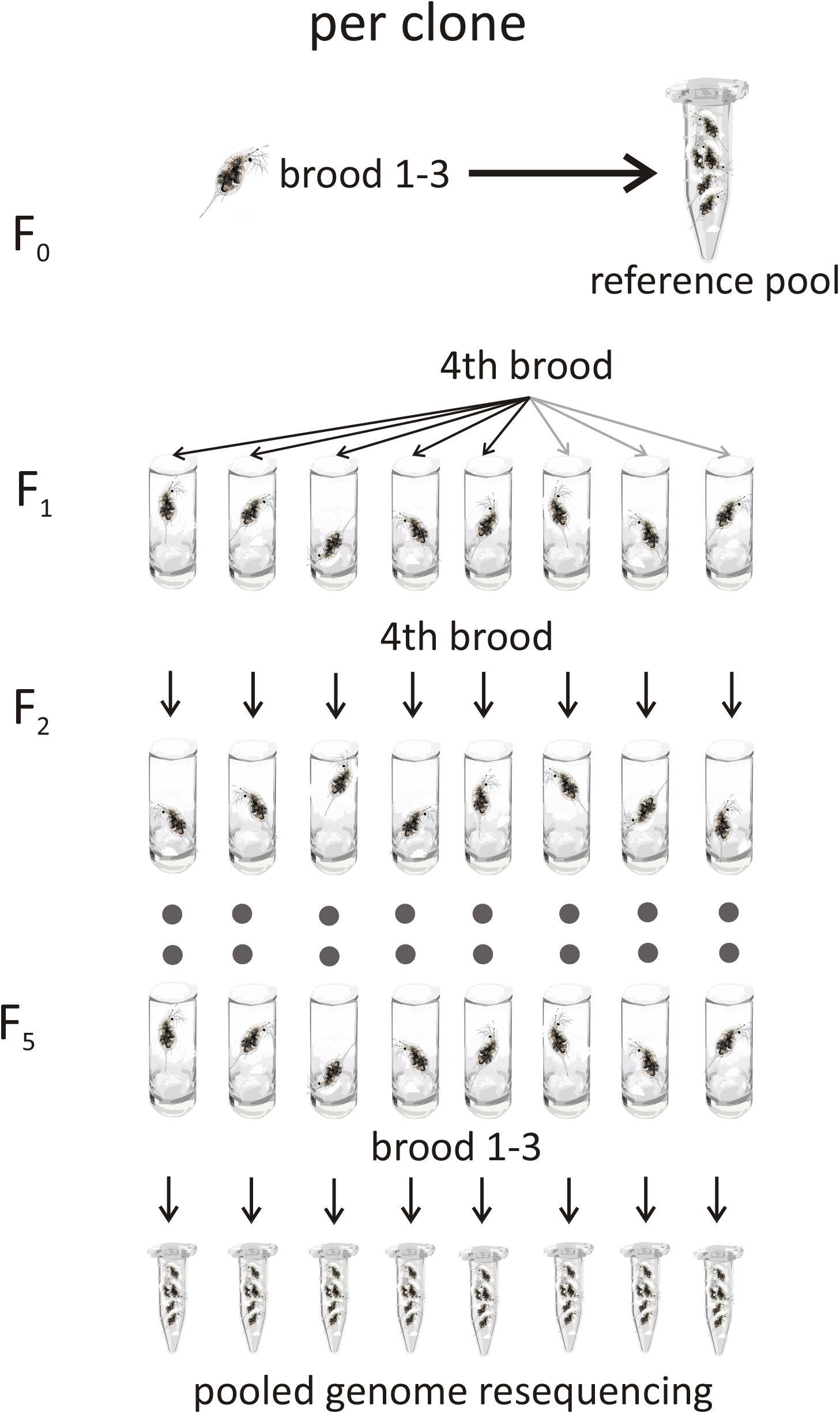
Schematic experimental set-up for the short term mutation accumulation lines per clone.

### Whole genome sequencing and bioinformatic processing

DNA was extracted for each pool of individual following a modified CTAB protocol, including an RNase step. The ancestor reference pool of each clone was sequenced to an expected mean coverage of 60X. After propagation for five generations broods of each of the MA-line was whole-genome sequenced to an expected mean coverage of 30X on an Illumina PE150 platform. Sequencing libraries were generated using NEBNext^®^ DNA Library Prep Kit following manufacturer’s recommendations. The genomic DNA was randomly fragmented to a size of 350bp by shearing, then DNA fragments were end polished, A-tailed, and ligated with the NEBNext adapter for Illumina sequencing, and further PCR enriched with P5 and indexed P7 oligos. The PCR products were purified (AMPure XP system) and resulting libraries were analyzed for size distribution by Agilent 2100 Bioanalyzer and quantified using real-time PCR. Reads were individually adapter clipped and quality trimmed, using Trimmomatic (Bolger, et al. 2014). Data was made available at ENA (acc. nos. ERS4993274-ERS4993294).

The cleaned reads of the ancestors and the MA lines were processed according to the best practices of the GATK-pipeline (McKenna, et al. 2010). Reads were mapped with bwa mem (Li and Durbin 2009) against the reference genome draft (Nickel, et al. in prep.) and filtered using a combination of Picard tools v1.123 (https://broadinstitute.github.io/picard/) to mark the duplicates and GATK v.3.3.0 (McKenna, et al. 2010) for realignment around indels and recalibration of bases. The resulting bam files were then prepared according to the input needed for accuMUlate (Winter, et al. 2018). Which means that each sample was individually identified at the sample (SM:) field of the read-group tag and merged together with Picard’s MergeSamFiles.

AccuMUlate was then run for each of the strains, J2 (1 ancestor and 7 MA lines), M5 (1 ancestor and 7 MA lines) and LC3.6 (1 ancestor and 4 MA lines) using the reference genome and specifying the following individual parameters for *D. galeata:* nucleotide frequencies of the reference genome (0.306 0.194 0.194 0.306, respectively), probability of sequencing error (0.001), ploidy of descendants (2) and ancestor (2).

The output table was then further filtered with a custom python script according to probability of a mutation/one mutation/of correct descendant genotype (>=0.90); number of reads matching the putatively-mutant allele in samples that are not mutants (=0); AD test statistic for mapping quality difference (>=1.95); p-value from a Fisher’s exact test for Strand Bias and Pair-mapping rate difference (>0.05). The final candidate list was then visually checked with IGV (Thorvaldsdóttir, et al. 2013) for validation.

To calculate the effective population size N_e_, we estimated Watterson’s theta (θ) (Watterson, 1975) based on a sample that consisted of 12 resequenced *D. galeata* genomes from Dobersdorfer See, Germany from Nickel, et al. (in prep.). We computed genotype likelihoods in ANGSD v0.931 (Korneliussen et al., 2014) from BAM files aligned to the reference genome for all 4-fold degenerate sites using the SAMtools model (option −GL 1). Sites were filtered for a minimum mapping quality score of 30, a minimum base quality score of 20 and reads that had multiple mapping best hits or secondary hits were removed. The folded site frequency spectrum was calculated with the realSFS program and used as prior to estimate per-site Watterson’s θ for all sites using thetaStat implemented in the ANGSD package (Korneliussen et al., 2014).

## Results

From the 24 MALs, 18 produced enough offspring in the fifth generation to isolate sufficient DNA for re-sequencing. The MAL were sequenced to an overall mean coverage depth of 34.64 (s.d. = 4.47, minmum mean coverage = 22.45, maximum mean coverage = 42.86). On average, 8.95 x 10^7^ sites (67% of the genome assembly, s.d. = 9.7 x 10^6^, min = 6.48 x 10^7^, max = 1.0 x 10^8^) were callable. In total, we scanned more than 1.6 billion diploid sites for mutations (Table 1).

**Table 1.**
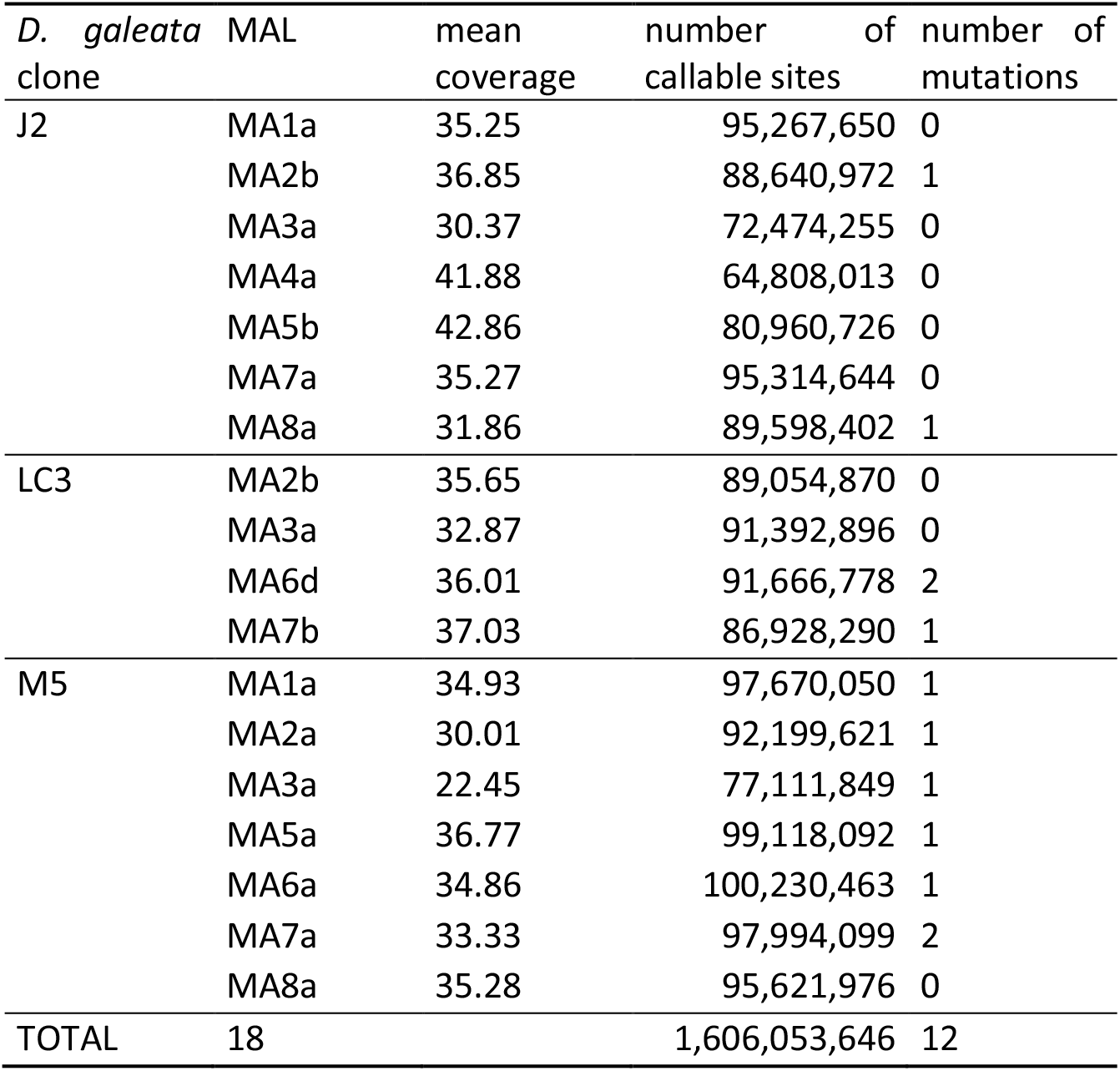
Information on the short term mutation accumulation lines (MAL) from three clones of *D. galeata* investigated.

**Table 2.** List of mutation positions, their characteristics and effect.

| *D. galeata*<br>clone | MAL | scaffold | position | SNM | transition<br>(ts) or<br>transversion<br>(tv) | amino acid<br>change | gene function an |
| --- | --- | --- | --- | --- | --- | --- | --- |
| J2 | MA8a | dgal52 | 163689 | G > A | ts | - | - |
|  | MA2b | dgal61 | 450819 | C > A | tv | - | - |
| LC3 | MA6d | dgal3 | 462655 | G > T | tv | - | - |
|  | MA7b | dgal98 | 307348 | A > C | tv | - | - |
|  | MA6d | dgal9 | 531976 | C > T | ts | - | - |
| M5 | MA5a | dgal9 | 390326 | G > A | ts | P > L | Cellular nucleic a<br>protein |
|  | MA1a | dgal24 | 857256 | A > C | tv | - | - |
|  | MA7a | dgal40 | 527171 | A > C | tv | - | - |
|  | MA3a | dgal57 | 335627 | C > G | tv | - | - |
|  | MA7a | dgal57 | 589433 | C > T | ts | - | - |
|  | MA2a | dgal121 | 469817 | A > C | tv | K > T | Density-regulate |
|  | MA6a | dgal270 | 8936 | G > T | tv | 4 fold<br>degenerate | unknown functio |

In the 18 MAL, we detected 12 single nucleotide mutations in 90 clonal generational passages (0.133 mutations per passage, Table 1). The rates among clones did not differ significantly (pairwise Poisson tests, p > 0.05 in all comparisons), therefore we report the mutation rate for all clones together. The haploid SNM rate μ was calculated as 0.745 x 10^-9^ (95% cf 0.39 x 10^-9^ – 1.26 x 10^-9^, Table 1). This rate was slightly lower than rates reported for *Daphnia pulex*, while all were substantially lower than the rate of *D. magna* (Figure 2). Using this rate, the mean θ estimate of 0.0092 and the relation θ = 4Neμ, the estimate for the long term effective population size was 3.09 x 10^6^ (95% cf 1.83 − 5.90 x 10^6^) for *D. galeata*.

**Figure 2.**
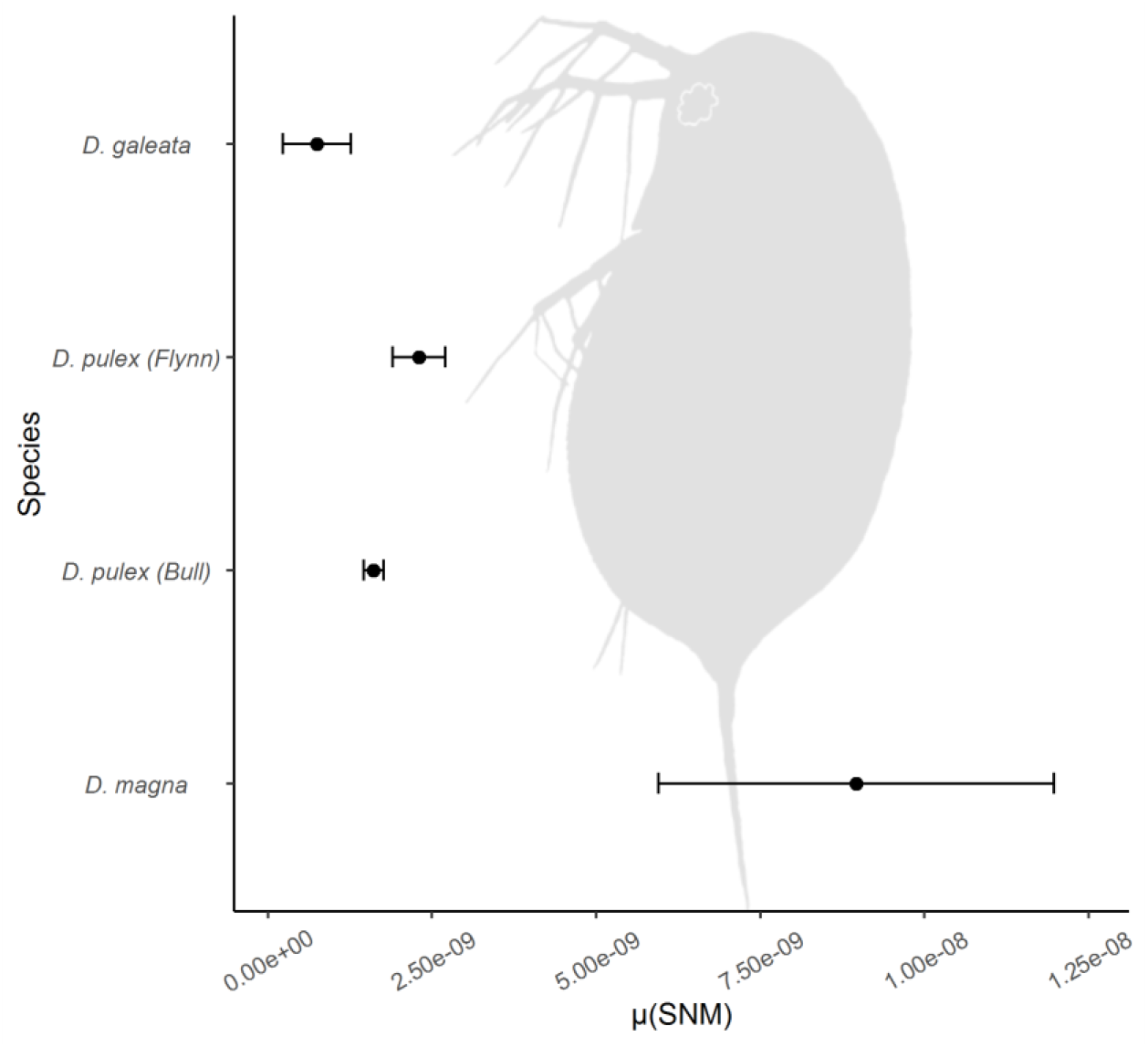
Haploid mutation rate (+/- 95% c.f.) of *Daphnia galeata* in comparison to other directly measured mutation rates of the genus.

Three of the twelve observed mutations (25%) were found in exons of predicted genes. This was within expectations given that the exon-space covers 22% and the gene-space 38.8% of the *Daphnia* genome assembly (Nickel et al. in prep.). Of the three exon located mutations, one (dgal270.8936) was a synonymous G >C change at a 4fold degenerate site in a protein of unknown function. The two others resulted in non-synonymous changes. The G > A change at dgal9.390326 in a gene annotated as *Cellular nucleic acid-binding protein* caused an amino acid change from Proline > Leucine. A gene annotated as *Density-regulated protein* showed an A > C transversion (dgal121.469817) that resulted in a Lysine > Threonine exchange.

The ratio between A/T > G/C and G/C > A/T mutations was 7/4 = 1.75, which is in line with the observed GC content of 38.75% in the *D. galeata* genome. The ratio of transitions (4) to transversions (8) was 0.5, which is exactly the unbiased expectation.

## Discussion

We report here for the first time a directly estimated clonal mutation rate for *Daphnia galeata*, a widely used model species. The obtained rate will significantly strengthen population- and comparative genomic approaches and serve as base line in evolutionary experiments of factors influencing the mutation rate. In contrast to other mutation rate estimates in *Daphnia* (Keith, et al. 2016; (Bull, et al. 2019; Flynn, et al. 2017; Ho, et al. 2020), which relied on MAL over several dozen generations, we have used the less time consuming short term mutation accumulation approach recently devised by (Oppold and Pfenninger 2017). While we obtained an accurate (low) mutation rate, the number of accumulated mutations was too low for meaningful analyses and comparisons of the mutational spectrum. However, information on the mutational spectrum will accumulate in future experiments to remedy this deficiency.

The spontaneous mutation rate of *D. galeata* reported here was slightly lower than rates estimated for *D. pulex* and much lower than for *D. magna.* Because the effective population size of the species was also the highest among the three species for *D. galeata* (N_e_ = 7.8 x 10^5^ in *D. pulex*, Lynch, et al. (2017) and 4.2 x 10^5^ in *D. magna*, Ho, et al. (2020)), our results support the drift-barrier hypothesis, which predicts that the mutation rate should be as low as drift limited selection permits, because mutations are generally deleterious (Sung et al 2012). We found only few mutations per clonal generational passage (0.133), indicating a remarkable replication fidelity at first sight. However, we measured here the mutation rate per clonal generation. Given that *Daphnia* clones go through several generations between sexual reproductions (Zaffagnini 1987), the cumulative mutation rate between sexual reproductions is likely at least a magnitude higher as the clonal mutation rate (and the calculated N_e_ accordingly lower), disregarding a potentially different mutation rate during sexual reproduction. Whether the use of a mutation rate estimate based on one parthenogenetic generation is appropriate to calculate the number of effectively sexually reproducing parents appears generally questionable. Lynch, et al. (2017) found a 2-5 fold discrepancy between N_e_ and the efficiency of selection in *D. pulex* compared to *Drosophila melanogaster*, which may have its cause in using the clonal mutation rate instead of the cumulative mutation rate between sexual reproductions.

Even though the number of mutations per clonal reproduction appeared to be low, this is put into a different perspective when considering the demography of the species. During peak densities, the number of individuals per square meter water column is in the order of 10^5^ - 10^6^ (Murtaugh 1985; Petersen 1983). Even small lakes (say, 1 ha) therefore harbour billions of individuals (10^9^-10^10^). Assuming that the mutation rate inferred here also applies to natural conditions, a fraction of 0.133 of them carries a single nucleotide mutation relative to the previous generation. Therefore, the demographic peak generation in the hypothetical lake alone carries 1.33 x 10^8^ – 1.33 x 10^9^ newly arisen mutations. With an estimated total genome size of about 1.6 x 10^8^, each genome position is therefore hit mathematically between 0.8 and 8 times by a mutation in such a population. Even if the density may be lower in other lakes and vary within lake, it is fair to assume that populations at least in moderately sized lakes are not mutation limited. Every possible mutation is practically always present in the population and in larger lakes, perhaps even in every clonal lineage. This almost permanent presence of exhaustive genetic variation should offer excellent opportunities for adaptive tracking of changing environmental conditions (Pfenninger and Foucault 2020), moreover since clonal reproduction should help to avoid stochastic loss of beneficial mutations (Kimura 1962; Messer and Petrov 2013). In addition, the occasional seasonal sexual reproduction allows to recombine favourable variation together. This extraordinary, mutation-driven propensity of *Daphnia* for rapid adaptation may be the background for the observed monopolisation of lakes by particular clones (De Meester, et al. 2002).

## Acknowledgements

The authors thank the LOEWE-Centre TBG funded by the Hessen State Ministry of Higher Education, Research and the Arts (HMWK).

## References

Acuna-Hidalgo R, Veltman JA, Hoischen A 2016. New insights into the generation and role of de novo mutations in health and disease. Genome Biology 17: 1–19.

Bolger AM, Lohse M, Usadel B 2014. Trimmomatic: a flexible trimmer for Illumina sequence data. Bioinformatics 30: 2114–2120.

Brede N, et al. 2009. The impact of human-made ecological changes on the genetic architecture of *Daphnia* species. Proceedings of the National Academy of Sciences 106: 4758–4763.

Bull JK, Flynn JM, Chain FJ, Cristescu ME 2019. Fitness and genomic consequences of chronic exposure to low levels of copper and nickel in *Daphnia pulex* mutation accumulation lines. G3: Genes, Genomes, Genetics 9: 61–71.

Charlesworth B 2009. Effective population size and patterns of molecular evolution and variation. Nature Reviews Genetics 10: 195–205.

Cui R, Kwak JI, An Y-J 2018. Comparative study of the sensitivity of *Daphnia galeata* and *Daphnia magna* to heavy metals. Ecotoxicology and Environmental Safety 162: 63–70.

De Meester L, Gómez A, Okamura B, Schwenk K 2002. The Monopolization Hypothesis and the dispersal–gene flow paradox in aquatic organisms. Acta Oecologica 23: 121–135.

Flynn JM, Chain FJ, Schoen DJ, Cristescu ME 2017. Spontaneous mutation accumulation in *Daphnia pulex* in selection-free vs. competitive environments. Molecular Biology and Evolution 34: 160–173.

Hall DJ 1964. An experimental approach to the dynamics of a natural population of *Daphnia galeata mendotae*. Ecology 45: 94–112.

Herrmann M, Ravindran SP, Schwenk K, Cordellier M (2018) Population transcriptomics in *Daphnia:* the role of thermal selection. Molecular Ecology 27: 387–402.

Hiroshi Akashi RMK, Eyre-Walker A 2012. Mutation pressure, natural selection, and the evolution of base composition in *Drosophila*. Mutation and Evolution 7: 49–60.

Ho EK, et al. 2020. High and highly variable spontaneous mutation rates in *Daphnia*. Molecular Biology and Evolution.

Ho SY 2014. The changing face of the molecular evolutionary clock. Trends in Ecology & Evolution 29: 496–503.

Huylmans AK, López Ezquerra A, Parsch J, Cordellier M 2016. De Novo Transcriptome Assembly and Sex-Biased Gene Expression in the Cyclical Parthenogenetic *Daphnia galeata*. Genome Biology and Evolution 8: 3120–3139. doi: 10.1093/gbe/evw221

Johnson T, Barton N 2005. Theoretical models of selection and mutation on quantitative traits. Philosophical Transactions of the Royal Society B: Biological Sciences 360: 1411–1425.

Keith N, et al. 2016. High mutational rates of large-scale duplication and deletion in *Daphnia pulex*. Genome Research 26: 60–69.

Kimura M 1962. On the probability of fixation of mutant genes in a population. Genetics 47: 713.

Klüttgen B, Dülmer U, Engels M, Ratte HT 1994. ADaM, an artificial freshwater for the culture of zooplankton. Water Research 28:743–746.

Korneliussen, TS, Albrechtsen, A, and Nielsen, R 2014. ANGSD: Analysis of Next Generation Sequencing Data. BMC Bioinformatics 15: 356.

Li H, Durbin R 2009. Fast and accurate short read alignment with Burrows-Wheeler Transform. Bioinformatics 25: 1754–1760.

Lynch M, et al. 2017. Population genomics of *Daphnia pulex*. Genetics 206: 315–332.

McKenna A, et al. 2010. The Genome Analysis Toolkit: a MapReduce framework for analyzing next-generation DNA sequencing data. Genome Research 20: 1297–1303.

Messer PW, Petrov DA 2013. Population genomics of rapid adaptation by soft selective sweeps. Trends in Ecology & Evolution 28: 659–669.

Miner BE, et al. 2012. Linking genes to communities and ecosystems: *Daphnia* as an ecogenomic model. Proceedings of the Royal Society Biological Sciences 279: 1873–1882. doi: 10.1098/rspb.2011.2404

Murtaugh PA 1985. Vertical distributions of zooplankton and population dynamics of *Daphnia* in a meromictic lake. Hydrobiologia 123: 47–57.

Oppold AM, Pfenninger M 2017. Direct estimation of the spontaneous mutation rate by short-term mutation accumulation lines in *Chironomus riparius*. Evolution Letters 1: 86–92.

Petersen F 1983. Population dynamics and production of *Daphnia galeata* (Crustacea, Cladocera) in Lake Esrom. Ecography 6: 285–294.

Pfenninger M, Foucault Q 2020. Quantifying the selection regime in a natural *Chironomus riparius* population. bioRxiv doi.org/10.1101/2020.06.16.154054

Pfenninger M, et al. 2015. Unique evolutionary trajectories in repeated adaptation to hydrogen sulphide-toxic habitats of a neotropical fish *(Poecilia mexicana)*. Molecular Ecology 24: 5446–5459.

Schiffels S, Durbin R 2014. Inferring human population size and separation history from multiple genome sequences. Nature Genetics 46: 919–925.

Sniegowski PD, Gerrish PJ 2010. Beneficial mutations and the dynamics of adaptation in asexual populations. Philosophical Transactions of the Royal Society B: Biological Sciences 365: 1255–1263.

Sung W, Ackerman MS, Miller SF, Doak TG, Lynch M 2012. Drift and the evolution of mutation rates. Proceedings of the National Academy of Sciences 109: 18488–18492

Tams V, Lüneburg J, Seddar L, Detampel J-P, Cordellier M 2018. Intraspecific phenotypic variation in life history traits of *Daphnia galeata* populations in response to fish kairomones. PeerJ 6: e5746.

Thorvaldsdóttir H, Robinson JT, Mesirov JP 2013. Integrative Genomics Viewer (IGV): high-performance genomics data visualization and exploration. Briefings in Bioinformatics 14: 178–192.

Weber A, Declerck S 1997. Phenotypic plasticity of *Daphnia* life history traits in response to predator kairomones: genetic variability and evolutionary potential. Hydrobiologia 360: 89–99.

Winter DJ, et al. 2018. accuMUlate: A mutation caller designed for mutation accumulation experiments. Bioinformatics 34: 2659–2660.

Zaffagnini F. 1987. Reproduction in Daphnia. In: Peters RH, de Bernardi R, editors. Daphnia: Mem. dell’istituto Ital. di Idrobiologia: Consiglio Nazionale delle Richerche. p. 245–284.

Zhang C, Jansen M, De Meester L, Stoks R 2019. Rapid evolution in response to warming does not affect the toxicity of a pollutant: Insights from experimental evolution in heated mesocosms. Evolutionary Applications 12: 977–988.

